# Identification and functional assessment of GPCRs across human adipogenesis

**DOI:** 10.64898/2026.07.22.739785

**Authors:** Jie Wu, Murray Polkinghorne, Eunice Liew, Iona Davies, Lili Davies, George Young, Ivan Andrew, Laurence Game, Katharine Lazarus, Patricia Ortega, Ahmed. R. Ahmed, David Carling, Tricia Tan, Ben Jones, Alice Pollard

## Abstract

Clinical obesity, defined as the presence of excess adiposity in conjunction with the presence of at least one clinical presentation of disease, remains a significant social and economic burden. Pharmacological weight loss agents based on glucagon-like peptide-1 receptor (GLP-1R) targeting are effective but exhibit significant side-effects leading to cessation of treatment, weight regain and, importantly, re-development of co-morbidities. Adipose tissue, now established as a central mediator of energy balance, endocrine signalling and inflammation, plays a significant role in the protection against metabolic disease onset. Loss of adipose tissue expandability, insulin sensitivity and function is thought to be a pivotal event in the transition to clinical obesity. Targeting adipose tissue dysfunction prior to or following development of metabolic disease remains a key therapeutic strategy either in conjunction with incretin-based weight loss therapy, or as a stand-alone therapy. The recent confirmation of functional glucose-dependent insulinotropic polypeptide receptor (GIPR) in mature adipocytes has led to a significant shift in our mechanistic understanding of dual GLP-1R/GIPR agonists such as tirzepatide, with the addition of adipocyte-specific targeting thought to underpin its clinical superiority to GLP-1R agonism alone. However, the expression, regulation and mechanistic targets of GIPR, and indeed many G protein-coupled receptors (GPCRs), in human adipocytes remains unclear, with adipocyte development being particularly under-studied in this regard. Here we use unbiased transcriptomic analyses of human adipocyte development with high temporal resolution to identify the onset of human GPCR expression, uncovering a previously undocumented surge in expression following adipogenic induction and elegant gene ‘waves’ throughout adipocyte development that may provide attractive pharmacological targets for adipose tissue dysfunction with or without weight loss. We functionally characterise GIPR, GLP-1 and CALCR/RAMP (Amylin) receptor activation at key differentiation timepoints, and identify a novel amylin response in early adipogenesis, presenting committed adipogenic precursors as a primary target of amylin signalling.

**Highlights:**

- Comprehensive transcriptomic analysis of human adipogenesis with improved temporal resolution and depth
- Identification and classification of >150 GPCRs differentially regulated across adipocyte differentiation
- Functional validation highlighting targeting of adipose stem cells and adipocytes using clinically approved receptor agonists
- Generation of novel human adipocyte stem cell lines from healthy and obese individuals to drive early target validation and exploration of adipocyte biology with increased pre-clinical power.

**Summary:** Comprehensive temporal RNA sequencing across human adipocyte development with a focus on G-protein coupled receptor expression and activity.

## Background

The burgeoning prevalence of obesity is a major public health challenge. Obesity-related ill health spans many disease areas, including classically “metabolic” conditions such as type 2 diabetes and metabolic dysfunction-associated steatotic liver disease (MASLD), but also “non-metabolic” manifestations including mobility problems and increased risk of certain cancers^1^. Given the importance of adipose tissue in physiological metabolic signalling, there is increasing recognition of the need to direct treatment towards “clinical obesity”, rather than excess weight per se^1–3^. Adipose tissue (AT), comprised of adipocytes, adipose stem cells, immune and vascular cells^4,5^, plays a pivotal role in maintaining energy homeostasis, balancing nutritional status and energy demand, integrating autocrine and endocrine signalling, regulating circulating hormone levels and protecting vital organs from lipid deposition^2,6,7^. The pathological transformation of AT, or adipose tissue dysfunction (ATDF), associated with metabolic disease onset is characterised by loss of adipocyte function, insulin insensitivity and pro-inflammatory cytokine production, leading to immune cell infiltration, fibrosis and stem cell senescence^8^. Adipose stem cells (ASCs) residing in the AT vasculature^3,9^ and reticular interstitium^10^ maintain adipocyte turnover through adipogenesis, contribute to signal integration, and provide a supporting network of extracellular matrix associations to maintain tissue health and structure. Adipogenesis is a finely tuned, coordinated response to a variety of stimuli (e.g., insulin, glucocorticoids), with dynamic gene expression facilitating changes to cell identity, morphology and function^11^. Perturbations in adipogenic trajectory at any stage can alter terminal cell function, promoting inflammation, senescence and fibrosis as well as impacting interactions with non-adipocyte cell types and driving ATDF. Identifying therapeutic targets to preserve and promote healthy adipogenesis at each stage of differentiation is an exciting strategy for the prevention of metabolic disease onset.

G protein-coupled receptors (GPCRs) form the largest family of protein targets for approved and investigational drugs. Since the development of glucagon-like peptide-1 receptor (GLP-1R) agonists such as semaglutide, there has been great interest in therapeutic targeting of gut hormone receptors to treat obesity. Most research has focussed on their anorectic actions, but many gut hormones have broader metabolic effects that could enhance or modulate their suitability for treating obesity and its clinical manifestations. Tirzepatide, a dual agonist for the GLP-1R and glucose dependent insulinotropic polypeptide receptor (GIPR), exerts direct adipocyte GIPR-mediated effects that may contribute to its overall metabolic benefits, such as the augmentation of post-prandial insulin sensitivity and stimulation of lipolysis during fasting^12^. However, little is known about whether tirzepatide or other GIPR-targeting agents act on adipocyte precursors or only mature adipocytes. The potential for adipocyte-related actions of other prime GPCR obesity targets, such as amylin^13^ and neuropeptide Y^14^ receptors, have been hinted at in other contexts but not formally investigated with therapeutic potential in mind.

Here we provide a transcriptomic map of human adipocyte development with improved temporal resolution, using patient-derived adipose stem cell lines to identify key regulatory networks governing GPCR expression. We functionally validated a selection of commonly targeted receptors using both naturally occurring and pharmacological ligands, revealing new potential therapeutic targets in adipocyte development. Our study identifies committed preadipocytes as targets of GPCR agonism; future work will identify down-stream consequences of long term agonism on adipose tissue depot turnover and function.

## Methods

### Reagents

Dulbecco’s Modified Eagle Medium/Nutrient mixture F12 (DMEM/F12) (Gibco) sodium pyruvate (Gibco), penicillin-streptomycin (Gibco), TrypLE express enzyme (Gibco), 3-(N-morpholino)-propanesulfonic acid SDS running buffer (Invitrogen), 4-12% Bis-Tris gels (Invitrogen), TRIzol (Invitrogen), LipidTOX green neutral lipid stain (Invitrogen), skimmed milk powder (Oxoid), BSA protein standard (Pierce), PageRuler Plus HCl (Honeywell Fluka), BSA Powder, methanol and pre-stained protein ladder were obtained from ThermoFisher Scientific (Loughborough, UK). All peptides were obtained from Wuxi Apptec (China). cAMP Dynamic HTRF reagents were obtained from Revvity (Rhondda Cynon Taff, UK).

### Human Tissue Ethics

Human adipose tissue samples used in this research project are collected and managed through the Imperial College Healthcare Tissue Bank (ICHTB). ICHTB is supported by the National Institute for Health Research (NIHR) Biomedical Research Centre based at Imperial College Healthcare NHS Trust and Imperial College London. ICHTB is approved by Wales REC3 to release human material for research (17/WA/0161). Informed consent was obtained prior to surgery, performed at St Mary’s Hospital London or biopsy, performed at Imperial College Clinical Research Facility (ICRF), with provision of a Tissue Bank Ethics Patient Information Sheet, cover letter and written consent given on day of procedure. All human samples were collected, handled and stored according to the Human Tissue Authority (HTA) UK codes of practice, in accordance with the Human Tissue Act 2004. Tissue and primary cell cultures were collected and stored under the subcollection ICS_AP_24_054, project R22001 at the Medical Research Council Laboratory of Medical Sciences, London UK.

### Primary human (healthy and obese) adipose tissue-derived stem cell isolation and culture

### Generation of healthy human immortalised (hTERT) adipose-derived stem cells (h-iADSC)

Healthy human immortalised adipose-derived stem cells (h-iADSC) used for sequencing in this study were obtained through an on-going collaboration with AstraZeneca and are subject to a material transfer agreement (MTA). The methods used to generate this resource are outlined in (51). Briefly, primary SVF isolated from a healthy female (30yrs) subject that had undergone population doublings were infected with a retrovirus containing the plasmid pBABEHygro-hTERT Retroviral Vector (Cell Biolabs; RTV-007), which expresses hTERT driven by a long-terminal-repeat promoter. The GP2-293 Packaging Cell Line which contains only the MoMuLV gag and pol genes, were transfected with pBABEpuro-hTERT Retroviral Vector using PolyJet DNA in vitro transfection reagent (SignaGen Laboratories, Rockville, MD). The viral envelope portion of the packaging function (env gene) was supplied by transiently co-transfecting pAmpho packaging vector (Clontech, 631530). Culture supernatants containing virus were collected at 24 h after transfection and filtered through a 0.45 μm filter. Primary SVF cells from human white fat at 80% confluence were infected with supernatants in the presence of 4 μg/ml Polybrene daily until cells reached 90% confluence. Cells were then treated with hygromycin (concentrations ranging from 400 μg/ml) in Endothelial Cell Growth Medium MV2 (Promocell). Once drug selection was finished, the cells were maintained in culture medium with 50 μg/ml hygromycin for 2 weeks. This viral work was covered by RA19-0044. Cells were maintained in Endothelial Cell Growth Medium MV2 for up to 15 passages. For all differentiation experiments, cells were transferred into DMEM/F12 with differentiation cocktail as described.

### Generation of patient-derived human immortalised (hTERT) adipose-derived stem cell lines (Healthy: AS_iSc 1, 2 and 3 Obese: h-iObSc-14, 15 and 16)

#### Healthy

Subcutaneous adipose tissue (abdominal) from three healthy volunteers (#1, 2, 3) was collected in saline under negative pressure via needle biopsy. **Obese:** Subcutaneous adipose tissue was removed from the peri-operative abdominal area of one female patient during laparoscopic bariatric surgery (#14), one male (#15) and one female (#16) via needle biopsy and washed in sterile PBS. Samples were transferred to DMEM/F12+1% Penicillin/Streptomycin, 10% Fetal Bovine Serum (FBS) for transport. Detailed patient information can be found in Figure S3.

Tissue was minced and digested in 1mg/mL Collagenase I, 1mg/mL Collagenase II and 2mg/mL Dispase II in serum-free DMEM/F12 for 35 minutes at 37oC with gentle agitation. The cell suspension was passed through a 100μm filter and centrifuged at 250g for 5 minutes. The pellet containing adipose stem cells (stromal-vascular fraction) was resuspended and plated in DMEM/F12, 10% FBS, penicillin/streptomycin supplemented with 4.5g/L glucose and sodium pyruvate into cell culture plates for expansion and treatment, or placed into a T75cm^2^ culture flask containing EGM MV2 (PromoCell), 10%FBS, 1% penicillin/streptomycin for subsequent immortalisation. Cells were cultured for two passages (P2) in EGM MV2, 10% FBS, 1% penicillin/streptomycin. A T75cm^2^ cell culture flask at 100% confluency was dissociated using 0.25% trypsin-EDTA and resuspended in 15ml EGM MV2, 10% FBS, 1% penicillin/streptomycin containing 4μg/ml polybrene for 15 min. Cells were infected with lentiviral vector LV[Exp]-EF1A>hTERT [NM_198253.3] (VectorBuilder, USA. Vector ID:VB900102-8315pav) for 24h. Cultures were maintained until passage 12, with primary SVF achieving 3 passages before loss of adipogenic capacity. Differentiation potential was examined by adipogenic index at P2, P6 (Figure S3C, D) and P12, with subsequent sequencing experiments performed at P6. Cells are denoted as human immortalised obese subcutaneous (h-iObSc) with patient ID (14-16), with healthy as AS_iSc_1-3.

### Adipogenic differentiation

At 70% confluency, adipogenic differentiation was induced using 1μg/mL Insulin, 3nM Triiodothyronine, 0.25μM dexamethasone, 0.5mM 3-isobutyl-1-methylxanthine and 0.2mM indomethacin, in DMEM/F12, 10% fetal bovine serum, 17.5mM glucose for 48h. For full adipogenic differentiation, cells were transferred into 1μg/mL Insulin, 3nM Triiodothyronine for a further 9 days.

### Immunofluorescence

Following differentiation, cells were briefly washed in PBS and fixed using 4% paraformaldehyde, 2% sucrose for 5 minutes at room temperature. Following fixation, cells were washed 3x in PBS and permeabilised using PBS, 0.1% Triton X100 for 5 minutes. Cells were washed 3x in PBS-0.1% fish skin gelatin and incubated in blocking buffer (PBS-0.1% fish skin gelatin, 10% normal goat serum) for 1h at room temperature. Cells were washed 3x in PBS and incubated for 30 minutes with LipidToxTM 488 neutral lipid stain and DAPI. Cells were washed 3x in PBS and imaged using an EVOS 7000 (ThermoFisher Scientific, UK). Well plate scanning was performed (automated) with autofocus performed once per well using all channels. Exposure was optimised on control wells with the lowest intensity in each channel and maintained throughout the scan.

### cAMP Assays

Cells underwent differentiation in 96-well plates as described above. For determination of cAMP responses to peptide stimulation, after removing culture media peptides resuspended in DMEM containing 0.1% casein and 1 mM IBMX were added to the wells and incubated for 10 minutes at 37°C. cAMP was determined using the cAMP Dynamic HTRF kit (Revvity) after lysing cells using the manufacturer’s lysis buffer. 3-parameter logistic fitting was performed using Prism. Responses were expressed as fractional change from baseline and then scaled using the mean Emax across all three differentiation days for each cell line/ligand.

### RNA extraction

Cells were lysed at indicated 24h timepoints in parallel from hiADSCs and h-iObSc-14 following adipogenic induction. Total RNA was isolated from cells by incubation with 1 ml TRIzol® (Life Technologies) per 1×10^6^ cells on ice. Samples were centrifuged at 10,000 x g for 15 minutes and the supernatant removed to a fresh tube. Chloroform (400 μl per ml TRIzol®) was added and the mixture centrifuged at 10,000 x g for 15 minutes at room temperature. The aqueous phase was transferred to an RNAse-free Eppendorf and absolute ethanol (0.53 x volume) added. RNA was purified using RNeasy Mini spin columns (Qiagen). RNA was eluted in 50 μl RNase free H2O and frozen at -80°C until required.

### RNA sequencing

RNA quality was assessed using the Agilent 2100 RNA 6000 Nano assay and libraries were prepared using the Watchmaker Genomics mRNA Library Preparation Kit with IDT xGen™ Stubby Adapter Unique Dual 8bp indexing, following manufacturer’s instructions. Library quality was evaluated using the Agilent 2100 High-Sensitivity DNA assay, and their concentrations measured using the Qubit™ dsDNA HS Assay Kit. Libraries were pooled in equimolar quantities and sequenced on a NextSeq 2000 to generate ∼ 50 million Paired End 60bp reads per sample.

### Bioinformatic Analysis

RNA-seq reads were processed using cutadapt v4.7^15^ to remove Illumina adapters, to quality trim at Q20, and to filter read pairs containing N bases or where either read <31 bps. Processed reads were quantified with Salmon v1.9.0^16^ using transcripts from the GRCh38 Ensembl v107 annotations. Salmon’s expectation maximisation procedures were set to enable modelling of sequencing and GC biases. Within R v4.4.1, transcript level counts from Salmon were imported and aggregated to the gene level using tximport v1.34.0^17^. Sample quality assessments were performed in relation to collected RNA integrity (RIN) values, sample concentrations used in library preparation (ng/ul), read GC content, and percentage read mapping. Four samples failing quality standards at this point were discarded. Normalisation, further PCA- and clustering-based quality control based on normalised values, and differential expression analyses were performed with DESeq2 v1.46.0^18^. Separately for healthy and obese samples, assessments of temporal changes were made using the likelihood ratio test (LRT), comparing models incorporating or excluding Timepoint as a factor (∼Timepoint vs ∼1) and WGCNA v1.73^19^ was used to cluster the rlog-transformed expression values of genes passing a q<0.01 cut off. Threshold powers for each group were determined following recommended practice from plots of scale free topology values for varying soft thresholding powers and final block-wise modules inferred from a signed network, allowing for genes to both increase and decrease in expression throughout the time course. In addition to the trajectory analysis, Wald tests were performed for specific timepoint samples between the healthy and obese groups. Independent hypothesis weighting was conducted to optimise the power of p-value filtering^20^ using IHW v1.34.0 and log2 fold change shrinkage was performed using ashr v2.2-63^21^ to reduce the impact of low expression on estimation of fold change values. Multiple correction testing adjustments were optimised by passing thresholds for both false discovery rate and log2 fold change at the point of results generation within DESeq2. Significance was assessed using a q value of 0.01. Gene ontology analysis was performed for clustered expression modules and for Wald test results (separately for increasing and decreasing genes within the comparison) using clusterProfiler v4.14.4^22^. Significance was assessed using a q value of 0.05.

### Statistics and Graphics

Heatmaps were created from genes differentially expressed (Wald test) relative to Day 0 (e.g., Day 1 vs Day 0) of genes passing the q<0.01 cut off, ranked by pAdj within each module, followed by log2fold-change ranking at either peak (for genes positively regulated), trough (for genes negatively regulated) or maximum +/- over Day 0 (for dynamic genes) during course of differentiation. Heatmaps, pie charts and graphs were created in GraphPad Prism v10.4.2. GPCR plots (tree and wheel plots) were created using GPCRdb^23^ data mapper with scales referring to gene cluster modules 0-6 (healthy) and 0-7 (obese). All GPCRs identified by LRT as differentially expressed across adipogenesis were included. Schematics were constructed using BioRender or Microsoft PowerPoint.

### Whole cell lysis and Protein extraction

Following differentiation, cells were washed briefly in ice-cold PBS and lysed rapidly in Radio-Immunoprecipitation Assay (RIPA) buffer, scraped and centrifuged at 13,500 rpm, 4°C for 10 minutes to pellet debris. The supernatant was collected for further analysis. Total protein content was determined by a detergent-compatible Bradford Assay (PierceTM) using a BSA standard curve.

### Western blotting

Unless otherwise stated, 10μg total protein was used for immunoblotting. Samples were denatured using 5x sample buffer containing 0.25M Tris-HCl pH 6.8, 0.25% bromophenol blue, 50% glycerol, 10% SDS, 0.5M DTT. Proteins were resolved by SDS-PAGE and transferred to a polyvinylidene difluoride membrane (Millipore Immobilon-FL) at 100 V for 90 minutes. Membranes were stained with Ponceau S to check protein transfer and blocked in 4% (w/v) bovine serum albumin (BSA) for 1 hour at room temperature. Unless stated otherwise, primary antibodies were diluted 1:1000 in TBS containing 4% BSA and 0.1% Tween-20, and incubated with the membrane for 4 hours at room temperature or overnight at 4°C. Membranes were washed extensively with TBS containing 0.1% Tween-20 before incubation with an appropriate IRDye secondary antibody (LI-COR Biosciences) in 4% milk powder-TBS 0.1% Tween-20 for 1 hour at room temperature. Blots were visualised using the Odyssey Imaging System (LI-COR Biosciences) and quantified using ImageStudio 5.2.

## Results

### Figure 1: A temporal transcriptomic map of human adipogenesis identifies gene expression waves associated with distinct phases of cell transformation

Several previous studies have contributed both transcriptome and proteome data during adipogenesis of various murine and human cell models, identifying both common pathways and those specific to cell type^11^. Importantly, the only comprehensive map of adipogenesis (>5 days profiled) in human cells failed to identify many receptor peptides, whether through low expression or technical limitations of mass spectrometry^11^. To enable accurate mapping of gene expression to inform functional receptor validation, we established a complete transcriptomic map of human adipogenesis using one healthy (h-iADSC) (AstraZeneca)^24^ and one obese adipose-derived stem cell line (h-iObSc-14) from subcutaneous adipose tissue (Figure 1A). We confirmed adipogenic potential and assessed adipogenic capacity through lipid accumulation (Figure 1B) and fatty acid synthase (FASN) protein expression (Figure S1A). We performed a differentiation time course in parallel with RNA extraction performed at 24h intervals from Day 0 (ADSC) to Day 11 (mature adipocytes), visualising trajectory by principal component analysis (PCA) (Figure 1C). Quality control excluded four samples based on low mapping rate and anomalous GC content (Figure S1 C-H). To establish gene expression patterns during adipocyte development we performed a Likelihood Ratio Test (LRT) analysis (Figure 1D) for both healthy and obese-derived cells, separating genes into modules (M#). Here we identified 7 (0-6) distinct patterns of expression in healthy and 8 (0-7) in obese-derived cells during differentiation. We visualised overlapping gene expression modules between healthy and obese-derived cell differentiation trajectories, highlighting commonality (e.g., hM2=OM1, 3525 overlapping genes) (Figure 1E).

**Figure 1:**
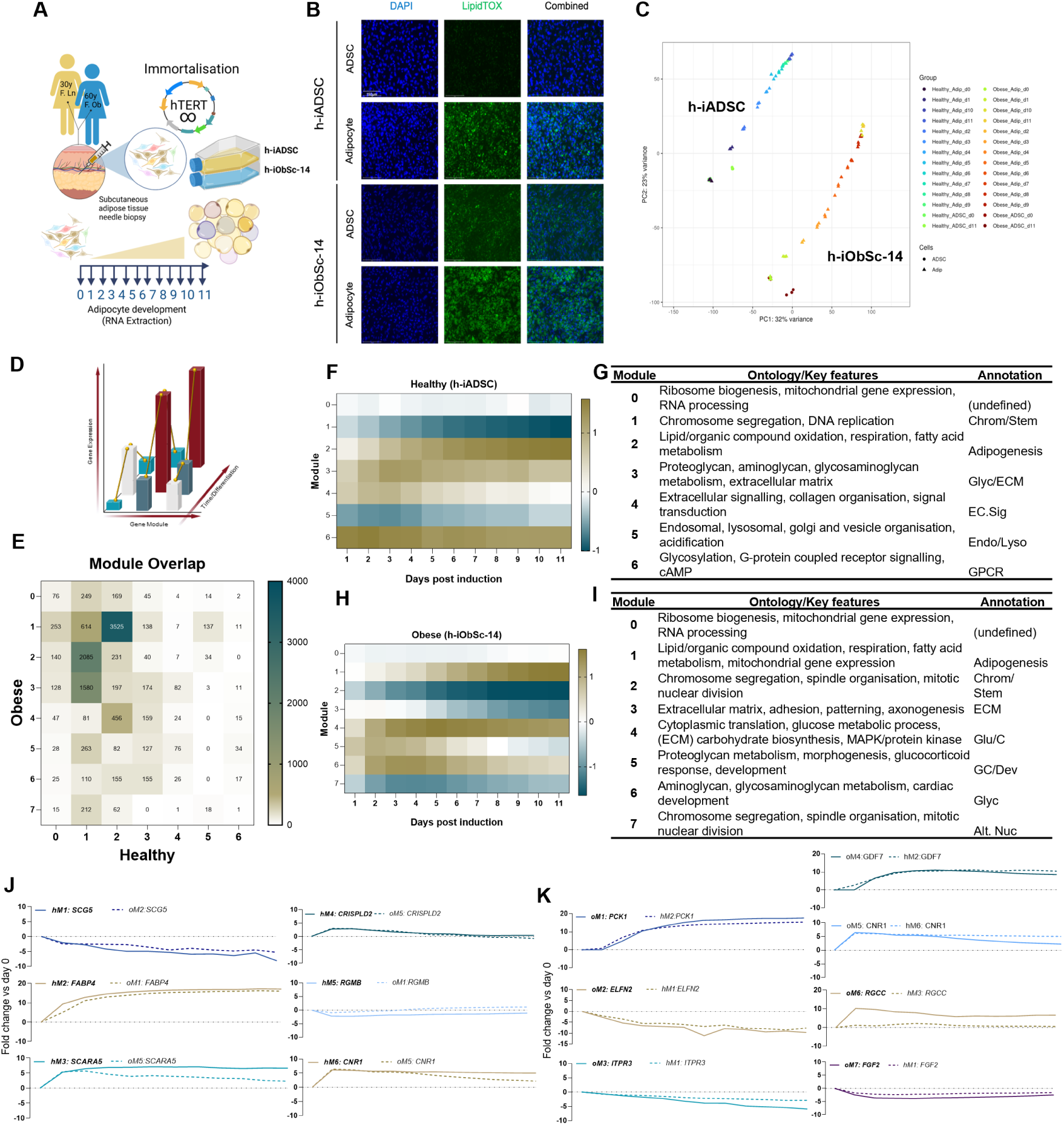
Adipogenesis is associated with dynamic regulation of key developmental, structural and metabolic pathways. **A)** Schematic depicting the generation of immortalised healthy (h-iADSC, AZ) and obese (h-iObSc-14) ADSC lines. **B)** Confirmation of adipogenesis in h-iADSC and h-iObSc-14 lines by lipid accumulation at day 11, prior to assessment of gene expression by RNA sequencing. Neutral lipid (Green, LipidTOX^TM^), nuclei (Blue, DAPI). **C)** Temporal RNA sequencing (24h time points) principal component analysis (PCA) **D)** LRT principle schematic, denoting change over time to provide only significant (pAdj<0.05) temporal shifts in gene regulation. Known phases of adipocyte development with proposed collection timepoints (Day 0-11) across adipogenesis. N=3 technical replicates per time-point. **E)** Overlap analysis identifying gene-module association between healthy and obese-derived cells. Data expressed in total gene count per overlap. **F)** Average change (Log2FC) in gene expression relative to day 0 aligned to modules 0, 1-6 (h-iADSC) and **G)** 0, 1-7 (h-iObSc-14). Significance at each timepoint determined by Wald test relative to day 0. Heatmap scale bars representative of average Log2Fold Change for ALL genes within module. **H)** Gene ontology allocation to each LRT module; h-iADSC (0-6) and **I)** (0-7) h-iObSc-14. Annotations for each module are given. **J)** Healthy and **K)** Obese expression module marker genes (ranked by pAdj then fold change). Marker gene expression and its module allocation for the alternate cell line are annotated by dotted line.

To characterise gene expression patterns during healthy human adipogenesis, we expanded and annotated each expression module identifying differentially expressed genes (Wald Test) relative to day 0. From this we obtained mean fold changes for all genes within each module (Figure 1F) and applied a gene ontology analysis to attribute biological function to each (Figure 1G). Using this analysis we identified several known expression patterns, including those known as ‘adipogenesis’ associated genes (hM2) with expression induced at day 2, following mitotic clonal expansion as well as downregulation of stem cell associated genes, annotated here as hM1, comprising chromosome segregation and DNA replication. We also identified several processes involved in early adipogenesis, including significant extracellular remodelling, proteoglycan and aminoglycan metabolism (hM3) peaking at day 3. Additionally, we observed a striking and dynamic regulation of gene subsets (hM4) relating to extracellular signalling and collagen organisation separate to those in hM3, following a distinct, rapid onset expression pattern with a peak at day 3. However, unlike those in hM4, these return to baseline in most instances, some with repression below that of day 0. Another inverse, dynamic regulation was observed in the organisation and acidification of organelles (hM5), negatively regulated during MCE and then restored in the following maturation phase indicating a considerable alteration to cellular shape, layout, and metabolism during adipogenesis. Finally, we identified rapid onset genes with peak expression obtained at adipogenic day 1 with persistent expression through to terminal cell differentiation, setting them apart from both hM3 in peak and hM4 in retention. hM0 denoted genes that did not fall into any other cluster. We observed that these genes followed a condensation pattern beginning with varied expression levels, converging throughout the time course, and returning to baseline variation at terminal cell differentiation. These genes were associated with RNA processing and ribosome biogenesis, in keeping with the initiation of multiple transcriptomic networks associated across differentiation.

To identify key differences between healthy and obese-derived ADSC differentiation trajectories, we applied the same LRT/Wald module analysis to the h-iObSc-14 data set (Figure 1H-I). Of the 7 gene-sets (oM1-7), we again found well-defined adipogenic, lipid and mitochondrial metabolism genes (oM1), with significant overlap in hM2, indicating successful commitment and adipocyte differentiation. We also found that, whilst well-defined in hM1, genes associated with chromosome organisation and MCE (downregulated over adipogenesis) were separated into oM2, 3, and 7 with distinct ontologies spanning mitotic division and chromosome segregation (oM2), ECM adhesion and patterning (oM3), and secondary chromosome segregation and nuclear mitosis (oM7) with a release of repression between day 4 and day 11. We identified oM4 as a sub-division of oM1/hM2 ‘adipogenic’ gene expression, genes associated with glucose and carbohydrate metabolism and MAPK signalling showing considerable disruption and divergence. In oM5 and 6, proteoglycan metabolism, glucocorticoid response and development overlapped with hM1, 2, 3, and 4, indicative of both disrupted and divergent signalling pathways with peak expression at day 2 followed by reduction, leaving expression only slightly elevated above baseline.

To further evaluate these gene clusters, we ranked all genes in each module by the fold change from peak/trough expression in that module (e.g., for hM5 the time-point with lowest expression). We then selected the top 20 by fold change and re-ranked these by adjusted p value (pAdj) according to LRT best fit to obtain module marker genes for healthy (Figure S2A-F) and obese (Figure S2G-M) gene expression modules. From these we selected the top 5 by pAdj to show adherence of individual genes to the overall trend within each module. Marker genes for each gene cluster (by fold change and pAdj) were then identified in both healthy and obese cells, displayed as overlays to show similarity or divergence between lines (Figure 1J, K).

Healthy hM1 marker gene chaperone protein secretogranin V (*SCG5*) expression decreased across adipogenesis similarly in both healthy (hM1) and obese (oM2) lines (Fig 1J). Additional hM1 marker genes included ECM-associated matrix metalloproteinase 1 (*MMP1*), BMP2/4 inhibitory protein gremlin 2 (*GREM2*), keratin associated protein 1-5 (*KRTAP1*-*5*), and leucine rich repeat containing 15 (*LRRC15*) (FigS2A). hM2, exhibiting a step-wise increase peaking at day 11, was marked by canonical adipocyte-associated gene fatty acid binding protein (*FABP*)4 (Fig 1J) in both lines (hM2/oM1), with additional hM2 genes including phosphoenolpyruvate carboxykinase (*PCK*)1 (also known as *PEPCK*-*C*), glycerol 3-phosphate dehydrogenase 1 (*GPD1*), protein phosphatase 1 regulatory inhibitor subunit 1A (*PPP1R1A*), and trafficking regulator of *GLUT4/SLC2A4* (*TRARG1)*, lipoprotein lipase (*LPL*), lipid droplet associated proteins perilipin (PLIN)1 and cell death inducing DFFA like effector (*CIDE*)*C*, and thyroid hormone responsive (*THRSP*) genes, all well-established hallmarks of adipocyte development (Figure S2B). All top5 hM2 markers overlap with oM1 top 20. hM3 marker scavenger receptor class A member (*SCARA*)*5*, previously identified as a critical regulator of adipogenic commitment in mesenchymal stem cells^25^, was rapidly upregulated between day 0 and 1 in healthy cells was identified in oM3, attributed to rapid onset and glycan metabolism in both lines, indicating overlap in early transcriptional activation. In obese cells SCARA5 decreased following rapid onset, and therefore assigned to dynamic cluster oM5. Additional hM3 markers included LIM domain only (*LMO*)3 (also known as dopamine associated transcription factor 1), hypoxia-associated EGFR interactor RIPOR family member (*RIPOR*)3, cell-cycle associated zinc finger and BTB domain containing (*ZBTB*)16, and carbohydrate sulfotransferase (*CHST*)2 (Figure S2C). The first of two highly dynamic modules, hM4, was marked by expression of cysteine rich secretory protein LCCL domain containing (*CRISPLD*)2, a recently identified, insulin-sensitising adipokine^26,27^ associated with weight-loss^28^, overlapping tightly in oM5. Additional hM4 markers included polypeptide N-Acetylgalactosaminyltransferase (*GALNT*)15, growth arrest specific (*GAS*)1, tripartite motif containing (*TRIM*)38 and protein tyrosine kinase 7 (inactive) (Figure S2D). Gene cluster hM5, exhibiting rapid, transient downregulation, were marked by BMP-associated repulsive guidance molecule (*RGM*)B, or DRAGON, a regulator of BMP signalling-established regulatory pathways governing adipogenic commitment in adipose stem cells^29^. In obese cells, RGMB associated with the adipogenic trajectory (oM1) despite a small early decrease, with elevated expression in adipocytes relative to day 0. Additional hM5 markers included transmembrane carboxylic/amino acid transporter solute carrier family 7-member (*SLC7A*)1, ER/endosome associated coronin (*CORO*)1C, calcyclin binding protein (*CACYBP*), and WD repeat and SOCS Box containing (*WSB*)2 (Figure S2E). Genes aligned to hM6, with rapid onset, retained gene expression (day 1 peak), associated with GPCR signalling and glycosylation, were marked by early expression of the cannabinoid receptor (*CNR*)1 (CB1), an established glucocorticoid-inducible GPCR^30^ typically characterised in the central nervous system (CNS) but also expressed highly by adipose tissue, where it is suggested to play a role in lipolysis^31^. *CNR1* also identified oM5, again defined by rapid onset, though with decline in expression during maturation. Additional hM6 marker genes included protein reversionless 3-like (*REVL3*), heme oxygenase (*HMOX*)1, feline leukaemia virus subgroup C choline and putative heme transporter (*FLVCR*)2, and neuropeptide Y receptor (*NPYR*) type 2 (Figure S2F).

We then identified marker genes for modules 0-7 in obese cells and mapped their expression in healthy equivalents. Similar trajectories between healthy and obese cells were observed in in those genes associated with adipogenesis (hM2/oM1), where *PCK1* (Fig 1K), *FABP4*, *THRSP*, and *LPL*, alongside adiponectin (*ADIPOQ*), were identified as marker genes (FigS2G). oM2 marker genes extracellular leucine rich repeat and fibronectin type III domain containing (*ELFN*) 2 (hM1) (Figure 1K), *LRRC15, GREM2* and *SCG5* were identified and consistent between healthy and obese cells (Figure S2H). These data indicated consistency between key adipogenic pathways across cell models. oM3, associated with axonogenesis, ECM and adhesion was marked by inositol 1,4,5-trisphosphate receptor type 3 (*ITPR3*) downregulation (Figure 1K) and established regulators of ECM extracellular matrix protein (*ECM*)1, G protein Ras-related C3 botulinum toxin substrate (*RAC)* 2 (Figure S2I). oM4, with high hM2 adipogenesis overlap, was defined by expression of growth differentiation factor (*GDF*)7 (Fig1K). Key differences were in the delayed onset of expression and slight decline towards terminal differentiation not observed in healthy cells. Also marking oM4 were the neuropeptide Y receptor (*NPYR*) type 1, inhibin subunit beta (*INHB*) B, arginine vasopressin receptor (*AVPR*) 1A and tenomodulin (*TNMD*), all mapped to hM2 (adipogenesis). oM5, exhibiting rapid (day 1) onset followed by return to or below baseline by day 10, shared marker gene CNR with hM6. oM6 marker gene regulator of cell cycle (*RGCC*) was identified in hM3, though with significant alteration in expression, independent of consistency in canonical adipogenesis (hM2/oM1).

### Figure 2: GPCR expression follows diverse temporal patterning during human adipogenesis

Our analysis identified many GPCRs differentially regulated across adipogenesis, some well-established and others with no known role. To further investigate the onset, expression, and regulation of GPCRs during adipogenesis, we utilised the International Union of Basic and Clinical Pharmacology (IUPHAR) GPCR resource, containing 400 known GPCRs to annotate all genes with a TRUE/FALSE GPCR status, grouped by module. We identified 143 GPCRs differentially regulated during healthy adipogenesis (Figure 2A) and 160 in obese adipogenesis (Figure 2B). By module, all but hM5 contained at least one GPCR, indicative of dynamic and coordinated regulation of GPCR classes throughout differentiation. We then identified GPCRs within each module by total count and as a percentage of total genes. In healthy adipocytes we found most GPCRs within hM1 and 2 by count (Figure 2C) and in hM6 by percentage (Figure 2D), representing bidirectional regulation of GPCR gene expression during commitment to the adipogenic lineage and high concentration within rapid onset genes. In obese cells, by count (Figure 2E) the majority of GPCR regulation associated with adipogenic genes (oM1), however by percentage expression is less discrete with distribution across oM2 and 3 as well as a far greater concentration within oM5 relative to total count (Figure 2F). We then mapped all identified GPCRs by module to the GPCR data base to identify receptor class using the GPCRdb data mapping tool^23^, allowing visualisation of class enrichment. We identified all adrenergic, GABA and formylpeptide receptors as differentially expressed across adipogenesis, significant enrichment of the frizzled (*FZD*) class C receptors and prostenoid receptors (*PTG*) and differential regulation of both the calcitonin receptor (*CALCR*) and calcitonin-like receptor (*CALCRL*) (Figure 2G). We first selected enriched receptor families with known adipogenic association; adrenergic, prostenoid, frizzled, NPY and calcitonin, revealing considerable divergence in expression patterns. Adrenergic receptors *A1A, A2A* and *B, B1, 2* and *3* associated with hM2 adipogenesis, *A2C* with downregulated hM1 chromatin/stem gene patterns, and *A1B* with early onset hM3 ECM/glycan genes (Figure 2H).

**Figure 2:**
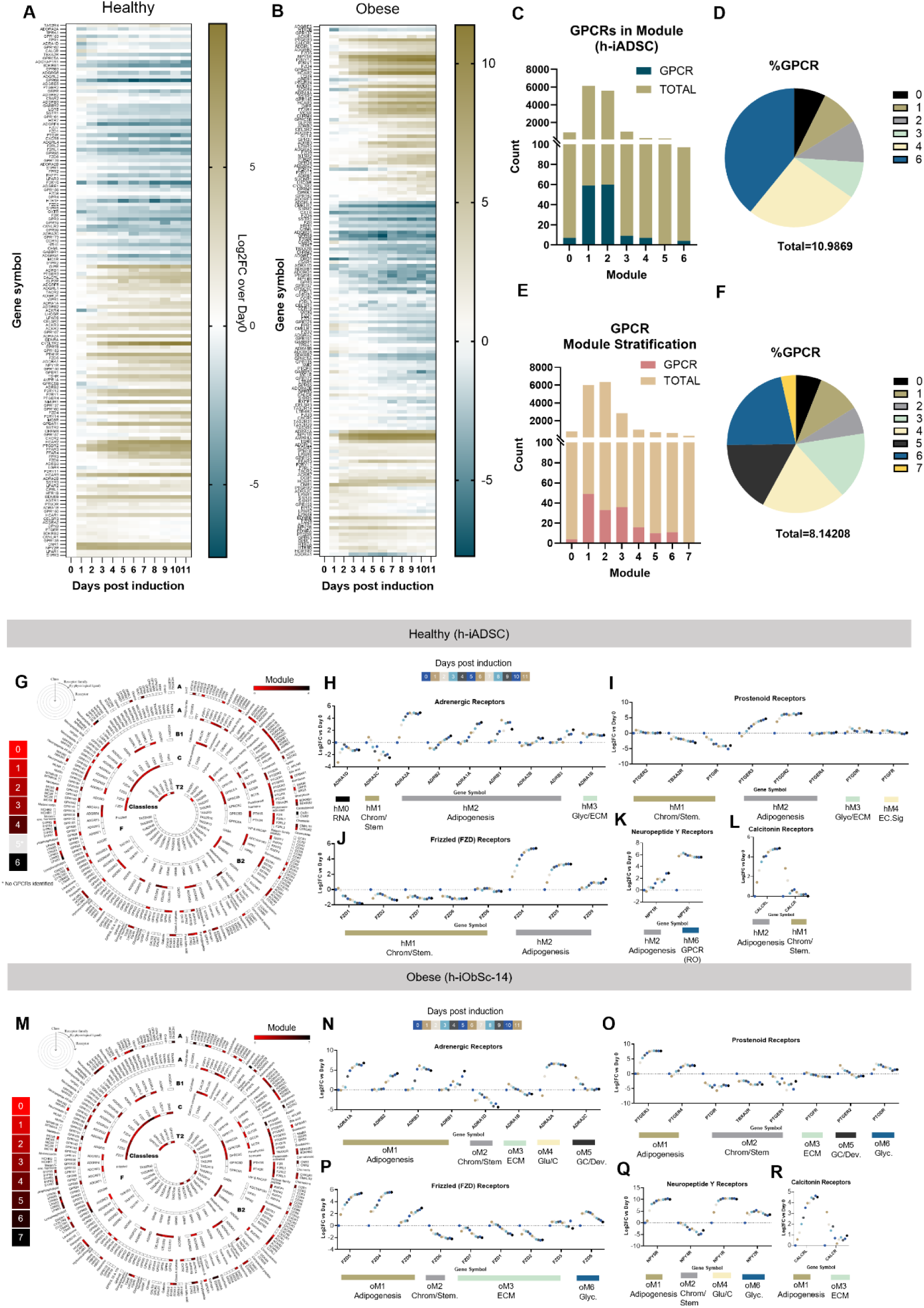
GPCRs are subject to dynamic regulation across adipogenesis. **A)** Heatmap displaying GPCRs identified by LRT as significantly changing during adipocyte differentiation, displayed by pAdj, in healthy h-iADSCs (143/400) and **B)** obese-derived h-iObSc-14 cells (160/400) **C)** GPCR count and **D)** % of total genes in module in h-iADSC (0-6) and (**E, F**) h-iObSc-14 (0-7). Total % GPCR in data set are shown. **G)** GPCR classification (GPCRdb) of identified GPCRs in h-iADSC. Modules denoted by scale bar. No GPCRs were identified in hM5. Gene expression (Log2FC over day 0) for selected classes of GPCRs; **H)** adrenergic, **I)** prostenoid, **J)** frizzled, **K)** neuropeptide and **L)** calcitonin receptors with known function in adipocyte development, annotated with module ID and identified GO term. **M)** GPCR classification and expansion of of identified GPCR classes **N-R)** in h-iObSc-14 adipocyte development modules. All data points are presented as Log2FC mean of n=3 technical replicates. Significance determined by LRT analysis q<0.01.

Prostaglandin receptors EP3 (*PTGER3)* and D2 (*PTGDR2)* aligned with adipogenesis, with contractile thromboxane receptor (*TBXA2R*) negatively regulated (Figure 2I). Low fold change, significant association was observed in *PTGER4* (hM2), and in *PTGDR* (hM3) and *PTGFR* (hM4). Frizzled receptor family members, well-established mediators of Wnt signalling, exhibited strong bi-directional association with downregulation of *FZD1, 2, 6, 7* and *8*, with *FZD4, 5* and *9* associating with adipogenic trajectory, peaking at terminal cell differentiation (day 9-11) (Figure 2J). Both neuropeptide Y receptors were upregulated during adipocyte development, with *NPY1R* following a canonical adipogenic trajectory and *NPY2R* exhibiting rapid onset expression at day 1 (Figure 2K). *CALCRL*, a component of the CGRP receptor when associated with receptor activity modifying protein (RAMP)1, and the adrenomedullin receptor (AM) when associated with RAMP2, followed an adipogenic trajectory. Expression of the *CALCR* previously identified in adipose tissue and adipocytes^32^, spiked at day 1 and then decreased below baseline throughout adipogenesis. We also identified proglucagon peptide family receptors *GLP2R* and *GIPR*, with no detected expression of *GLP1R*, confirming previous studies^12^. We performed this analysis for our obese cell line (Figure 2M), noting consistency in adrenergic receptor gene regulation with the exception of *ADRA2C*, downregulated during adipogenesis in healthy cells but following a transient upregulation in obese cells (Figure 2N). Divergence in *PTGER2* (oM5, GC/Dev) and notable absence of *PTGDR2* induction were observed in obese cells (Figure 2O). Frizzled receptor *FZD4, 5* and *9* (adipogenesis, oM1)*, and 6* (chromatin, stem oM2) expression patterns were conserved. *FZD1,2* and 7 were delayed in downregulation (day 3-4) and clustered with ECM signalling, possibly indicating slower transcriptional modification in this line, independent of adipogenic onset. We observed a transient upregulation of *FZD8* not observed in healthy cells (Figure 2P, Figure S2E). Most striking was the considerable induction of *NPY5R* in obese cells (Figure 2Q). *NPY5R* expression was considerable only in obese cells (Figure S3F), consistent with previous findings^33^. *CALCRL* and *CALCR* expression patterns were comparable, though with *CALCR* spike pronounced and aligning with oM3 (Figure 2R).

### Figure 3: Functional association of GPCRs differentially regulated across adipogenesis in healthy h-iADSCs

To further characterise GPCR expression across adipogenesis we generated GPCR tree plots annotating receptor class, ligand type, family, and gene ID for each gene expression module. This classification allowed visualisation of class and ligand enrichment or depletion at key adipocyte development timepoints without assumption of adipogenic association. Those genes downregulated during adipogenesis (hM1) (Figure 3A) and those conversely upregulated (hM2) (Figure 3B) exhibited high class diversity with notable switches including the exchange of lysophospholipid receptors (*SIPR1*, *2* and *5*) for free fatty acid receptors (*FFAR3* and *4*), an enrichment of *ADRA1A, 2A, 2B, ADRB1-3* in mature adipocytes, and loss of developmental *FZD1, 2, 6, 7* and *8* for lipid-associated *FZD4, 5* and *9*. Among hM1 (downregulated) were aGPCR adhesion molecules *ADGR L4, L2, G1*, *G6, F4, E1 E2, E5,* and *B3* (Figure 3A), previously reported in the context of adipogenesis, with hM2 containing *B2, D1, F5, L1*, and *CELSR2* as adipocyte-associated aGPCRs (Figure 3B). *GIPR* and *GLP2R* were significantly increased during adipogenesis. Those in hM3, with rapid induction and mature cell reduction (excluding *EDNRB*), were exclusively class A. A key feature of this analysis is the alignment of orphan receptors with unknown or poorly defined function to phases of cell development. Class A orphans *GPR 3, 4, 19, 39, 63, 68, 85, 161, 173, 176,* and *LGR5* were downregulated (Figure 3A), with *GPR78*, *146, 153, 160*, and *LGR4* associated with hM2 adipogenesis (Figure 3B). hM4 (transient, up) contained Class A bradykinin receptor (*BDKRB*) 2, chemokine receptor *CMKLR1* and two adhesion receptors *ADGRA2* and *CELSR3* (Figure 3D). No GPCRs were identified in hM5 (transient, down). Rapid onset GPCRs (hM6), in addition to those previously identified as markers (*CNR1, NPY2R*) were the lysophospholipid receptors sphingosine 1-phosphate receptor (*S1PR)3,* known to promote PPARγ/C/EBPα signalling^34^, and lysophosphatidic acid receptor (*LPAR) 1*, as yet uncharacterised in early adipocyte development.

**Figure 3:**
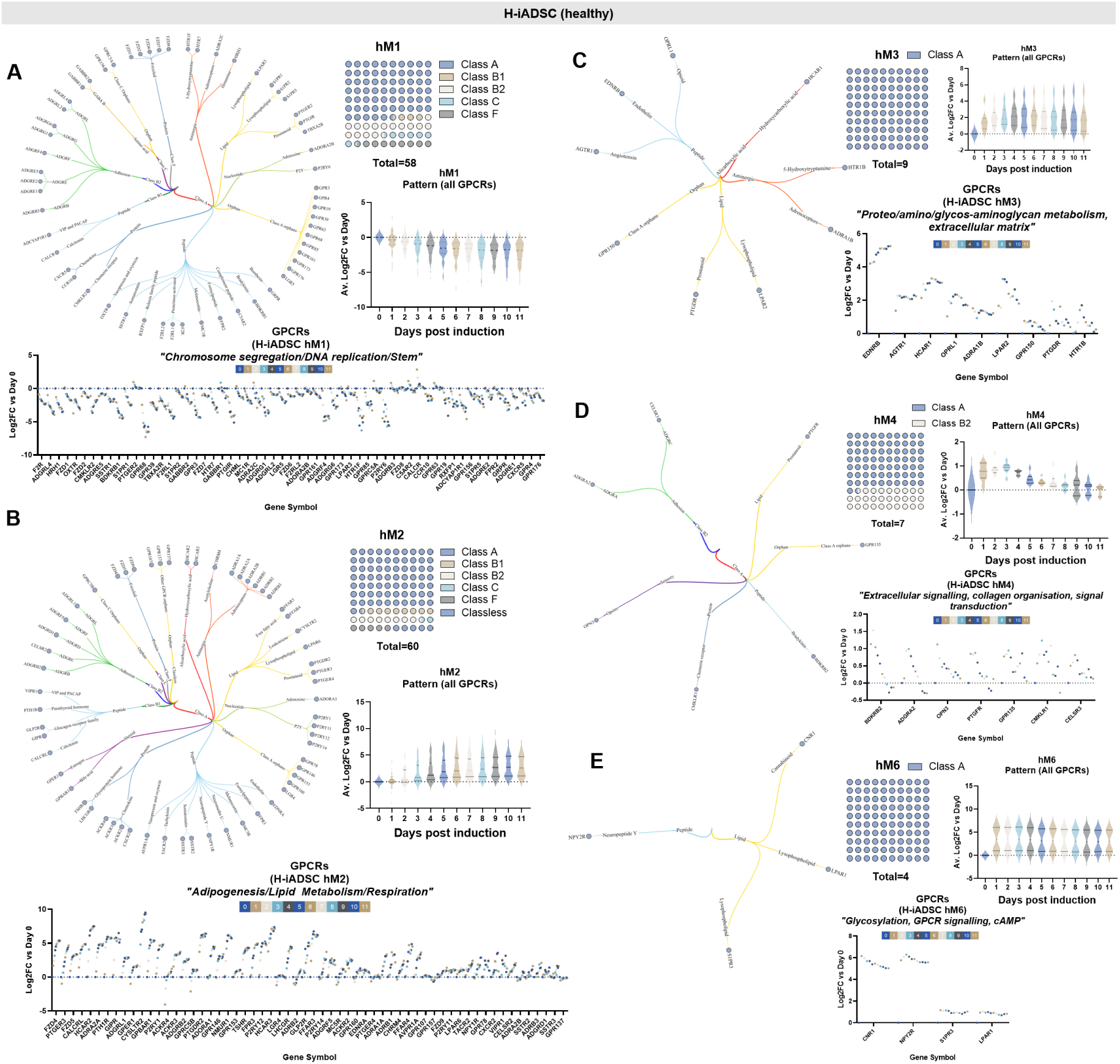
Functional association of GPCR gene expression modules in h-iADSC adipocyte development. GPCRs displayed from inner to outer edge as a tree plot to assign class, type, ligand and gene symbol. Class assignment is shown as 10×10 dot plots. Average of total GPCR expression Log2FC for each timepoint displayed as violin plot and each individual gene represented in dot plot, with each point an average of three technical replicates. **A)** GPCRs aligned to hM1, (chromatin segregation, DNA replication and stem cell). **B)** GPCRs aligned to hM2, (lipid metabolism, respiration, ‘adipogenesis). **C)** All GPCRs aligned to hM3 (proteo/amino/glycosaminoglycan metabolism, ECM). **D)** All GPCRs aligned to hM4 (EC-signalling, collagen organisation, signal transduction). **E)** All GPCRs aligned to hM6 (Glycosylation, GPCR signalling, cAMP). No GPCRs were identified in hM5.

### Figure 4: Functional association of GPCRs differentially regulated across adipogenesis in obese-derived-hiObSc-14 cells

Our data have established both h-iADSCs and h-iObSc-14s to be both accurate and comparative models of adipogenesis, with significant overlap in adipogenic gene expression induction (hM2/oM1), lipid accumulation (Figure 1A), and protein expression (Figure S1A). However, the extent to which each line can represent biological variation in all aspects, or to what extent these data may be extrapolated to other individuals, is unknown. We therefore approached this analysis independent of our previous findings, instead contrasting those falling into alternate modules to indicate divergence. For those re-assigned, or not identified by LRT, we consulted rLog gene expression data to obtain absolute value for comparative analysis. For stem/chromatin associated (downregulated during adipogenesis) module oM2 (hM1) (Figure 4A) we again identified *ADGRL4, G1, E2,* and *D1*, as well as *CELSR1* (Figure S2C). In contrast we observed reassignment of *ADGR L2* (oM4) (Figure 4D, S2A) and *G6* (oM3) (Figure 4C, S2D) as well as no significant expression change in *F4* (Figure S2B), *E1, E5,* and *B3*. *ADGRF4* (Figure S2B), rapidly downregulated in h-iADSCs, was detected at low levels in h-iObSc-14 and expression subsequently remained low, below threshold for LRT/Wald test identification. Similarly, *FZD6* (Figure S2E) was associated with canonical ‘stem’/downregulated genes, whereas *1, 2, 7,* and *8* were distributed between oM3 and oM6, in both cases exhibiting early increases in expression followed by down-regulation during maturation, with *FZD8* (oM6) the most pronounced. Orphan GPCR *GPR153*, positively regulated with healthy adipocyte development, repositioned in oM4 (previously shown to have high hM2 adipogenic overlap) (Figure 4D). RLog counts showed high basal expression of *GPR153* in both lines, though with differential regulation across adipogenesis confirming module reassignment (Figure S2G). *GIPR* and *CALCRL* again associated with adipogenesis (Figure 4B), however *CALCR* was reassigned to dynamic oM3 (Figure 4C), highlighting our previous identification of differential regulation.

**Figure 4:**
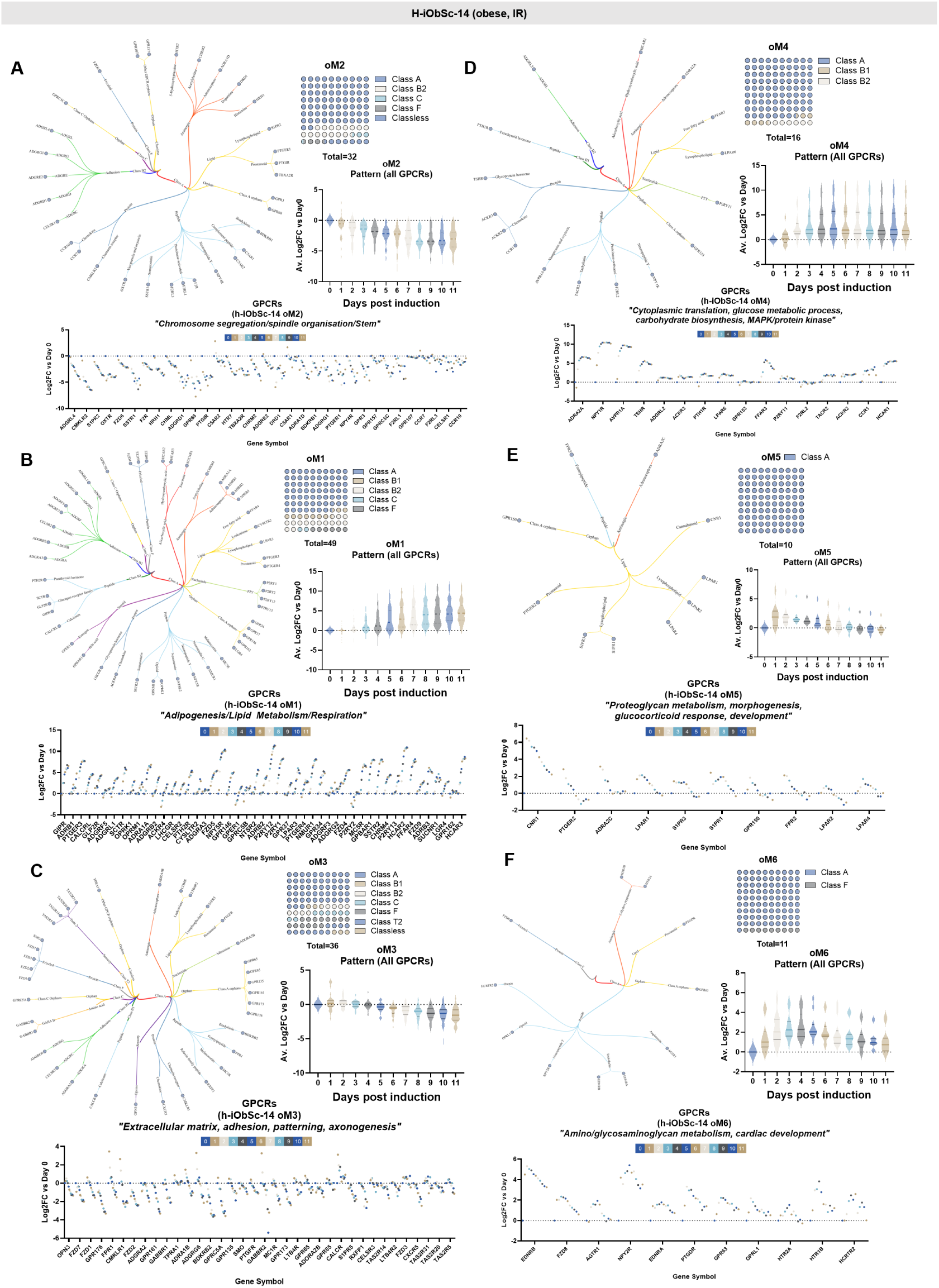
Functional association of GPCR gene expression modules in h-iObSc-14 adipocyte development. GPCRs displayed from inner to outer edge as a tree plot to assign class, type, ligand and gene symbol. Average of total GPCR expression Log2FC for each timepoint displayed as violin plot and each individual gene represented in dot plot, with each point an average of three technical replicates. **A)** GPCRs aligned to oM1, (lipid metabolism, respiration, ‘adipogenesis). **B)** GPCRs aligned to oM2, (chromatin segregation, DNA replication and stem cell). **C)** GPCRs aligned to hM3 (ECM, adhesion, patterning, axonogenesis). **D)** GPCRs aligned to oM (Cytoplasmic translation, glucose metabolic process, carbohydrate biosynthesis, MAPK/protein kinase) **E)** GPCRs aligned to oM5 (proteoglycan metabolism, morphogenesis, glucocorticoid response, development). **F)** GPCRs aligned to oM6 (amino/glycosaminoglycan metabolism, cardiac development).

### Figure 5: Functional characterisation of selected GPCRs identifies GIPR and AMYR as differentially responsive during adipogenesis

GPCRs are frequently expressed at low levels in cells and are difficult to detect via traditional methods for protein expression. We therefore selected candidate genes and used publicly available proteomic data sets (Klingelhuber *et al.,* 2023) to overlay our models with protein expression data from pre-existing human ADSC lines; Simpson-Golabi-Behmel syndrome (SGBS) cells, immortalised TERT-APCs similar to our own, primary hAPCs from the same donor, together with immortalised SVF-APCs (hWA)^35^ (Figure 5A). These data did not detect GIPR, GLP1R, NPY, or receptor activity-modifying protein (*RAMP)* family members in any line tested whether due to low expression or technical limitations of the mass spectrometry methods used. The data did, however, confirm expression of adipogenesis-associated GPCRs FZD4, PTGER3, and CALCRL as well as a transient regulation of hM3-ECM associated proteins endothelin receptor beta (EDNRB) and angiotensin II receptor type (AGTR) 1 (Figure 5B), confirming in part the ability of our data to predict GPCR protein expression. We then evaluated publicly available data through the AT Knowledge portal, and through recent human single-cell/single-nuclei adipose tissue studies, noting low coverage of most GPCRs. We identified *GIPR* expression in mature human adipocyte populations (Figure S5A), as well as enrichment in mural and lymphoid populations in data extracted from Emont *et al., 2022* and Miranda *et al.,* 2025 via Single Cell Portal. *CALCR* detection in adipocytes was low but highly enriched in myeloid cells (Figure S2B). *CALCRL* appeared enriched in all cell types identified, with highest expression in mature adipocytes and endothelial cells (Figure S5C). Additional data sets from the AT Knowledge portal failed to identify *GIPR* (Figure S5D) or *CALCR* (Figure S5E) in adipocytes, however CALCR was again enriched in mature adipocytes relative to other cell types. Both *GIPR* and *CALCRL* were identified as late adipogenic genes in trajectory analysis (Figure S5G, H), with disparity between CALCRL mRNA and protein expression observed by Klingelhuber *et al*., 2024^11^. We examined an additional study by Loureiro *et al.,* 2023 exploring transitioning adipocyte precursors and adipocytes. Neither GIPR nor CALCR were identified by this study, using two independent donors, however CALCRL was again enriched in adipocytes (Figure S5I). Given the recent indication in murine models, the expression of GIPR on human and murine adipocytes (Figure S5), and the increased interest in the targeting of GPCRs in metabolic disease, we sought to assess the functional presence of GIPR and amylin receptor components (CALCR/RAMPs) throughout adipogenesis, by measuring cAMP responses to appropriate endogenous ligands or pharmacological ligand derivatives. Here we used four healthy (h-iADSC, AS-iSc1, 2, 3) and three obese (h-iObSc-14, 15, 16) novel cell lines (Figure S4A-D) to add significant biological representation. Aligning with the temporal trend in *GIPR* expression, pronounced responses to GIP were detected at day 10 (d10) with smaller responses detected earlier in the differentiation time-course. The same pattern was observed with the dual GIPR/GLP-1R agonist peptide tirzepatide, which is presumed to be GIPR-mediated on account of the lack of detectable adipocyte *GLP1R* expression, absent functional response to GLP-1, and previous data^12^.. Recognising that *CALCR* expression showed a peak of expression at day 1 (d1)/day 2 (d2), it was noteworthy that cAMP responses to the amylin receptor family endogenous agonist ligands calcitonin and amylin, as well as those of the investigational anti-obesity amylin agonist cagrilintide, were also highest at d0 and d2 and had declined by d10. This is the first time, to our knowledge, that functional differences between pharmacological GPCR targeting ligand derivatives have been demonstrated in human adipocyte precursors and mature adipocytes. Our data has therefore identified and functionally validated significant alterations in GPCR gene expression in early human adipogenesis, with reproducible function across multiple individuals, presenting a novel therapeutic window to influence adipocyte development.

**Figure 5:**
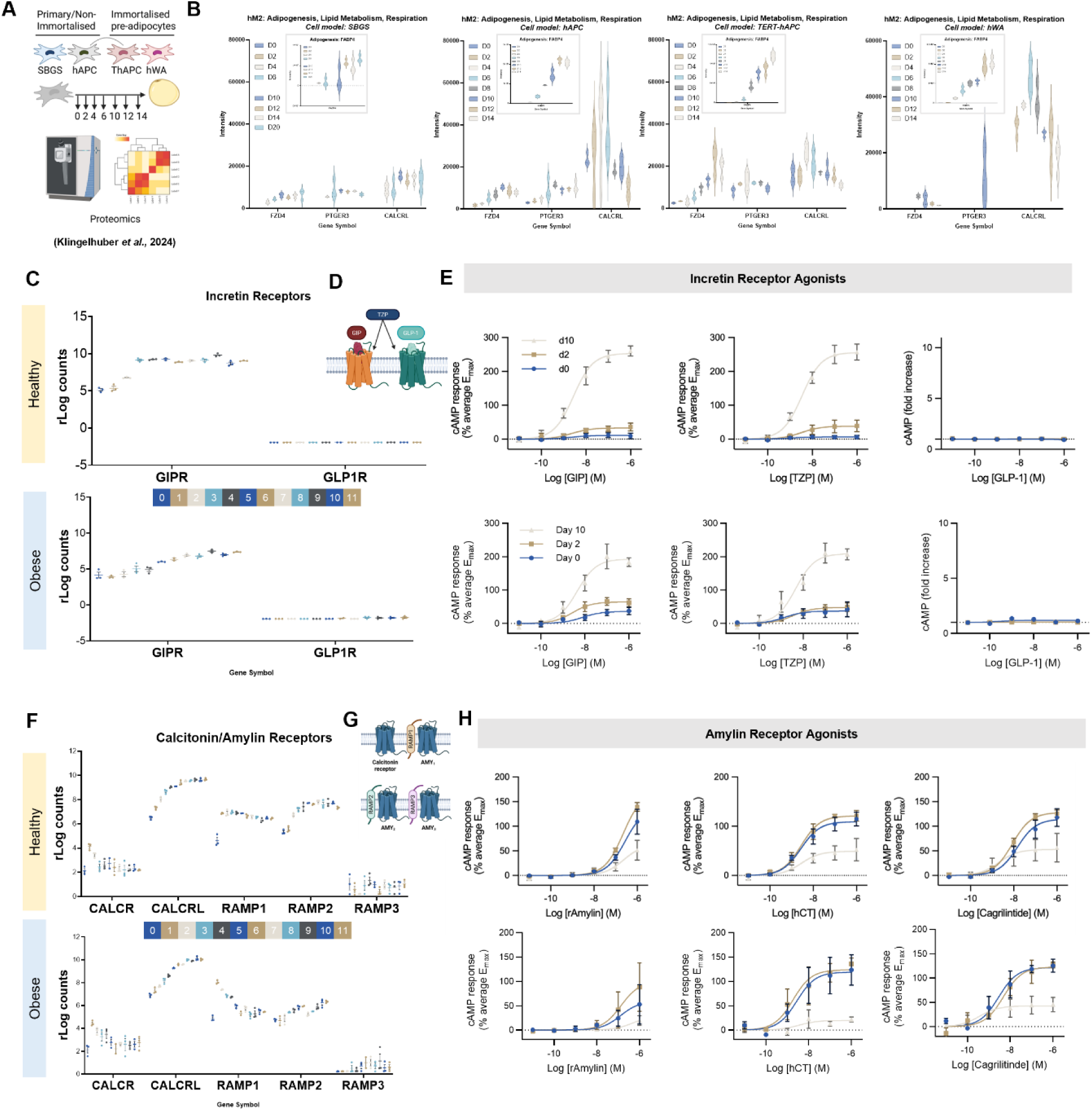
Functional assessment of GPCRs in human adipose-derived stem cells reveals opposing roles for GIPR and AMYR during adipogenesis. **A)** Publicly available proteomic data set (Klingelhuber *et al.,* 2024 doi: 10.1038/s42255-024-01025-8) methodology. Cells used include SBGS, hAPC (primary), hTERT-APC (immortalised hAPC, linked with hAPC), and hWA (immortalised lines). Data downloaded from ProteomeXchange under identifier PXD047412. **B)** Extracted data for all cell models with detected proteins in top5 genes aligned to hM2 to validate expression. Insets: fatty acid binding protein 4, control for adipogenesis. GIPR, GLP-1, CALCR, NPY and RAMP family not detected. **C)** *GIPR* and *GLP1R* (rLog) expression across human ADSC development. **D)** Mechanism of action of GLP-1, GIP and Tirzepatide (TZP). **E)** Dose-response curves following 10 minutes incretin treatment (cAMP production, calculated as fractional change from baseline and then scaled using the mean Emax for each time-point, except GLP-1 [fold-change from Emin]) at day 0 (undifferentiated ADSCs, day 2 (committed pre-adipocytes) and day 10 (mature adipocytes). **F)** Expression of calcitonin receptor (*CALCR*), calcitonin-like receptor (*CALCLR*) and *RAMPs* during adipogenesis. **G)** *CALCR* and *RAMP*-associated amylin receptor conformations for functional validation. **H)** Dose-response curves from 4 independent healthy donor cell lines (h-iADSC, AS_iSc-1-3) and 3 obese-derived cell lines (h-iObSc-14-16) following 10 minutes ligand treatment (cAMP production, calculated as in (E)) at day 0 (undifferentiated ADSCs, day 2 (committed pre-adipocytes) and day 10 (mature adipocytes). N=4(h), 3(ob) biological replicates (patient lines), with 4technical replicates per agonist concentration. hCT: human calcitonin, rAMY: rat amylin.

## Discussion

The success of GLP-1 receptor agonists for obesity treatment has led to great interest in developing new anorectic agents as adjuncts to GLP-1 for greater weight loss. There is recognition, however, that obesity treatment should target the clinical manifestations of obesity and not only excess weight *per se*. Given the centrality of adipose tissue function to metabolic health in the context of obesity, targeting receptors enriched in early ADSC differentiation, including GPCRs, is a promising therapeutic strategy for the improvement of adipose tissue expandability, particularly with regard to insulin sensitisation, reducing pro-inflammatory and pro-fibrotic stem cell trajectories, and promoting healthy adipocyte differentiation, independent of weight loss. Though powerful, many single-cell/nuclei and proteomic studies lack the depth to identify low-abundance proteins, and current single-cell studies achieve only partial coverage biased to highly expressed genes^4,5,11,36^. GPCRs are often poorly detected and, as we have demonstrated, are often absent from both single-nuclei and proteomics data sets examining adipogenic gene/protein expression. Our study combines improved temporal RNA-sequencing, identifying ∼150 GPCRs differentially expressed during human adipogenesis, with functional GPCR characterisation using plate-based cAMP assays. Kaczmarek *et al.,* 2023 approached a similar question using the RNA sequencing database FATTLAS, containing RNA signatures of obese and non-obese whole adipose tissue^37^. Four GPCRs were taken forward in a murine pre-adipocyte model 3T3-L1, all of which were identified in our study also. However, this study highlights, using 3T3-L1 and subsequently HEK293T for functional validation, that the study of GPCR signalling in primary human adipose tissue-derived cells is challenging^37^. Here we have focused on those GPCRs commonly targeted therapeutically, with commercially available ligands, with increased biological power through our novel cell line generation. Our data provide a scalable human resource for the evaluation of GPCR function in adipocyte biology. Using these data, we have demonstrated direct effects of GIP and amylin on mature adipocytes or differentiating ADSCs from four healthy and three obese donor cell lines, providing valuable biological evidence for therapeutic targeting.

Amylin analogues such as cagrilintide unquestionably drive weight loss via appetite suppression, but our data raise the possibility that effects on adipocyte development could also influence their metabolic phenotype, especially given the peak of *CALCR* expression early in our differentiation time-course. Direct effects of amylin on lipogenesis in mouse fibroblast-derived 3T3-L1 adipocyte-like cells have been described, as has STAT3 signalling induction in mature human adipocytes^32,38^, but, to our knowledge, the presence of functional amylin receptors during human ADSC differentiation has not been reported previously. Higher expression of *RAMP1* and *RAMP2*, compared to *RAMP3*, throughout adipocyte development indicates that *AMY1R* and *AMY2R* are the dominant amylin receptor subtypes, which could be relevant to adipocyte/ADSC targeting by emerging amylin analogues that in some cases show differences in their selectivity for each receptor subtype. For example, eloralintide selectively engages AMY1R over AMY3R when compared to cagrilintide, and preferentially leads to loss of fat mass over lean mass^39^. Future work will explore the role of amylin signalling in early adipogenesis and potential use of amylin analogues as a modulator of adipocyte stem cell function. We additionally identified *CALCRL,* which when partnered with RAMP1, forms a receptor for CGRP, and with RAMP2/3 forms the adrenomedullin (AM) receptor.

Expression of GIPR in adipocytes has long been debated, with our data in line with recent reports unambiguously confirming its presence^12^. Effects of GIPR agonism on mature adipocyte function have gained traction as mediators of the metabolic actions of tirzepatide^12^, especially as the anorectic effects of GIPR agonism that are seen in preclinical pharmacotherapeutic studies are not recapitulated in humans. However, little attention has been paid to possible effects of GIP on adipocyte precursors. One study reported the presence of *GIPR* mRNA during differentiation of SGBS preadipocytes, discussing its early onset, but did not confirm this functionally in differentiating cells^40^. Our time-course data both confirms previous findings and highlights the presence of functional GIPRs early in adipocyte development. GIPR signalling has also been shown to improve insulin sensitivity (phospho-Akt) and glucose uptake (2-DG) in mature adipocytes^12^. Though progress has been made in the study of insulin sensitivity in cultured adipocytes via manipulation of culture conditions favouring glucose uptake^41^, these do not address naturally occurring biological variability between donors or the role of insulin sensitisation in early adipogenic induction. Future work will focus on the insulin-signalling modulation of GIPR and amylin agonism in adipocyte commitment and differentiation across biological replicates.

Our analysis identified known and novel transcriptional patterning, including dramatic gene induction at poorly explored in other studies focused on day 2 and 10. Many of these genes, aligned to hM3 and 4 as well as oM5 and 6, are transient and their functions in adipogenesis less well explored than those associated with fatty acid metabolism or the loss of stem- and Wnt-associated gene expression. Previous studies using a proteomic approach have uncovered similar programs; Klingelhuber *et al.,* identified a similar patterning termed ‘transient’, where enrichment of proteins associated with lysosomes and glycosidases were dynamically regulated, and performed subsequent characterisation of organelles across adipogenesis, identifying shifts in abundance, localisation, and function. The association of orphan receptors with potential functional consequences will hopefully provide a starting point for further investigation in adipocyte development and in wider fields. The addition of our patient-derived data implies consistency across models, providing evidence in additional human cell lines with a wider age range, and identifying organelle biology during adipogenic commitment as a significant event. Future work will undoubtedly explore this in the context of lysosomal disorders, nutrient signalling and dietary interventions, where regulation of mitochondrial and lysosomal turnover and function are of clinical importance. Our findings add significant depth to these previous studies, with the addition of human cell lines to further functionally validate all remaining orphan receptors in future work.

## Conclusion

Here we provide a comprehensive, improved temporal analysis of human adipogenesis, identifying and classifying GPCRs associated with transcriptional patterning to provide underpinning data for pharmacological exploration of novel GPCR-targeting drug candidates. We identify, for the first time, functional differences between GIPR and amylin receptor activity in human adipocytes and ADSCs/committed APCs respectively, the latter previously unknown to the field. Given the increased interest in targeting adipose tissue dysfunction independent of weight loss, identification of GPCRs expressed by ADSC/APC populations and adipocytes independent of each other provides an exciting therapeutic strategy for harnessing the adipose tissue niche, not just canonical adipocyte targeting. Many adipose tissue disorders, including congenital, familial, and acquired lipodystrophies as well as those associated with life cycle (menopause, ageing) and hormonal imbalances are resistant to weight-loss intervention or independent of total fat mass. These, in conjunction with emerging data identifying those non-responsive to incretin-based weight-loss therapies, represent a significant unmet clinical need. Our data and cell lines therefore provide a valuable resource for further studies and will invite early pre-clinical investigation into the use of combined GPCR agonist approaches to target human adipocyte development, reducing reliance on murine cell models for functional validation.

### Limitations

Our study utilises human immortalised ADSC lines, 6 developed in house and one through collaboration (n=7), suitable for *in vitro* exploration of adipose tissue development, a significant improvement on existing resources. These include two male and five female subjects. However, as we, and others, have demonstrated, both intra- and inter-individual heterogeneity make the evaluation of pharmaceuticals challenging. Future work will increase sex- and age-matching in cell line use to explore and control for sex-differences that are well documented in adipose tissue. Similarly, our cell lines are derived from the abdominal subcutaneous adipose tissue; given the known differences between depots our future work will include visceral and organ-associated adipose tissue cell lines currently under development, and those from alternate subcutaneous regions. Our study is conducted in monolayer culture and cannot accurately recapitulate cell-cell interactions in the 3D adipose tissue environment, nor do they capture the multi-cellular, dynamic environment of AT, lacking immune cell and endothelial populations critical for tissue function, a feature not yet incorporated by the field in a multi-omic-compatible format capable of high-throughput analyses such as these. We and others are actively improving these systems to better recapitulate both tissue and population heterogeneity. Lastly, we appreciate that not all GPCRs can be profiled functionally in a single study and have selected those with commercially available FDA approved ligands for our functional validation, instead focussing on demonstrating efficacy across biological replicates. Future studies, both our own and across the field, will utilise these data and a growing repertoire of cell lines to validate functionality, downstream signalling and potential intervention strategies in future work.

## Author Contributions

**A.P, B.J, D.C and J.W** conceived the study. **A.P** generated h-iObSc-14-16 and AS_iSc1-3 cell lines. **J.W, M.P, E.L, I.D, L.D, B.J** and **A.P** designed and performed experimental procedures. **T.T** and **A.R.A** provided and supported access to human tissue samples. **P.O** performed surgical collection of adipose tissue samples. **K.L** performed needle biopsies to obtain tissue samples. **I.A** and **L.G** performed library preparation and RNA sequencing. **G.Y** performed bioinformatic analysis of the data set. **J.W, M.P, I.D, B.J** and **A.P** performed data analysis and presentation. **A.P** and **B.J** wrote the original draft manuscript. All authors were involved in the writing, editing and final drafting of the manuscript.

## Funding Acknowledgements

**J.W** was supported by the Imperial College London MRes programme. **M.P** is supported by a technical research and innovation specialist position within Imperial College London. **E.L** is supported by a technical research and innovation specialist position within Imperial College London. **L.D** was supported by an Imperial College London undergraduate studentship. **I.D** is supported by a research and innovation specialist postdoctoral position within Imperial College London. **K.L** is a Diabetes UK Sir George Alberti Research Training Fellow (grant reference number 23/0006515). **D.C** is supported by an MRC grant MC-A654-5QB10. **T.T** is supported by the NIHR BRC and grants from MRC, Diabetes UK and NIHR. **B.J** acknowledges funding support from the Medical Research Council (MR/Y00132X/1 and MR/X021467/1), the Wellcome Trust (301619/Z/23/Z), Diabetes UK, the Eli Lilly LRAP programme, and Metsera Inc. **A.P** was funded by a BBSRC Discovery Fellowship (BB/W009633/1) and Imperial College London Institute of Clinical Sciences.

## Acknowledgements

h-iADSCs were kindly provided by AstraZeneca as part of an ongoing MTA. The authors acknowledge the access to human tissue provided by the Imperial College NHS Trust and Imperial College Human Tissue Bank (ICHTB), and for their support of this work. The authors thank the patients and volunteers who contributed samples for this study and the Imperial College Clinical Research Facility (ICRF).

## Data and resource availability

All data generated in this study will be deposited into GSEA for public accession upon publication. H-iADSCs (AstraZeneca) are subject to MTA. All cell lines generated in this study are currently available through collaboration.

