## Supplemental Figures for "Identification and functional assessment of GPCRs across human adipogenesis"

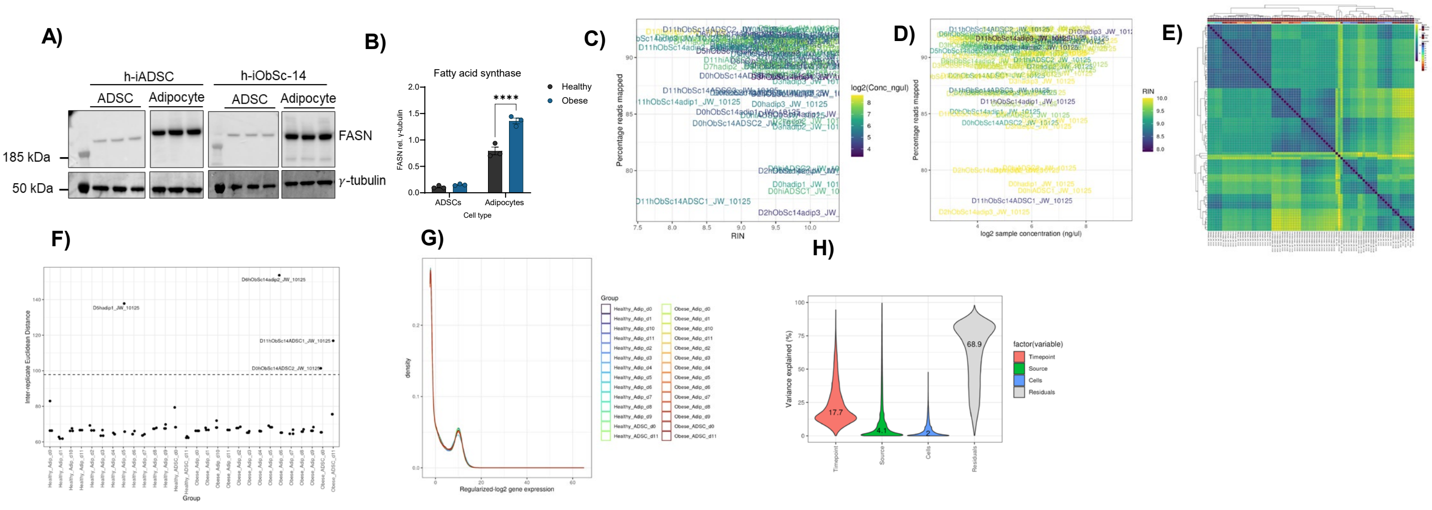

**Supplementary Figure 1: DESeq2 RNA sequencing QC for h-iADSC and h-iObSc-14**

**A)** Fatty acid synthase (FASN) protein expression and **B)** quantification of western blot, identifying adipogenesis independent of imaging/RNA. **C)** RNA Integrity scoring for all submitted samples. No samples were excluded based on RIN (>7.5). **D) Read mapping** across sample concentration. **E)** Inter-sample distance **F)** outlier detection identification of four samples with anomalous GC content and combined lower mapping rate. Two of four identified were ADSC controls (D0hiObSc-14 (ADSC2) and D11hi-ObSc-14 (ADSC1)), and the other two technical replicates for hiADSC adipogenic Day 5 and h-iObSc14 Day 6. These samples were subsequently excluded from the analysis. **G)** Regularised log2 gene expression and **H)** source of variation, attributing highest variation by timepoint, then source (healthy vs obese), then cells (ADSC vs Adipocyte).

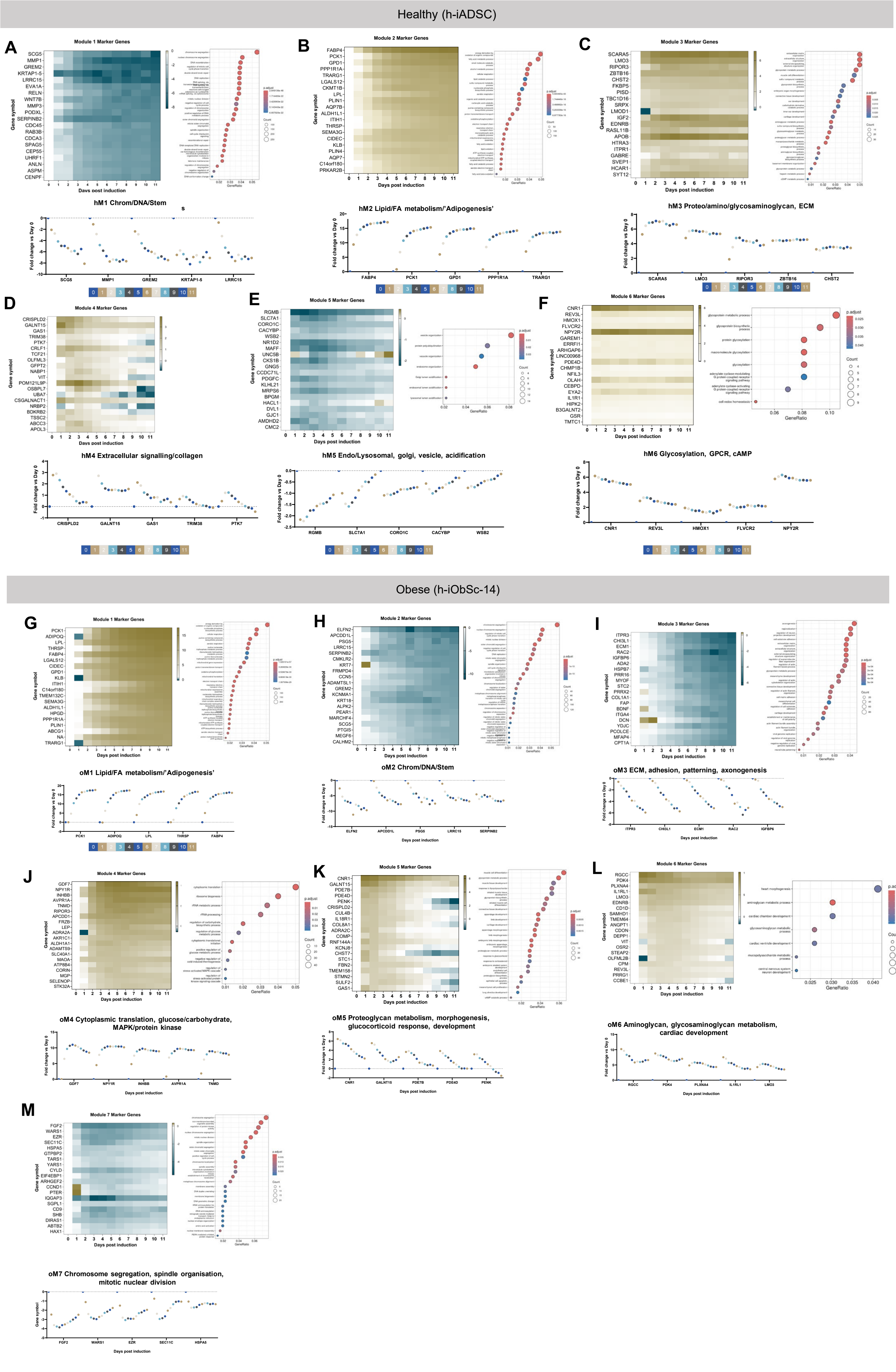


**Supplementary Figure 2: Marker gene and gene ontology analysis of identified gene modules in healthy and obese cell lines. A-F) Healthy h-iADSC and G-M) Obese h-iObSc** Marker gene identification per module; top 20 by pAdj (LRT-Wald test Log2FC relative to day 0) displayed by heatmap. ClusterProfiler Gene Ontology dot plots for all genes within each module are shown. Top 5 marker gene expression by pAdj, highlighting patterning and dynamic range within each module. hM4 did not provide dotplot, gene ontology performed using DAVID Functional Annotation tool GO_BP.

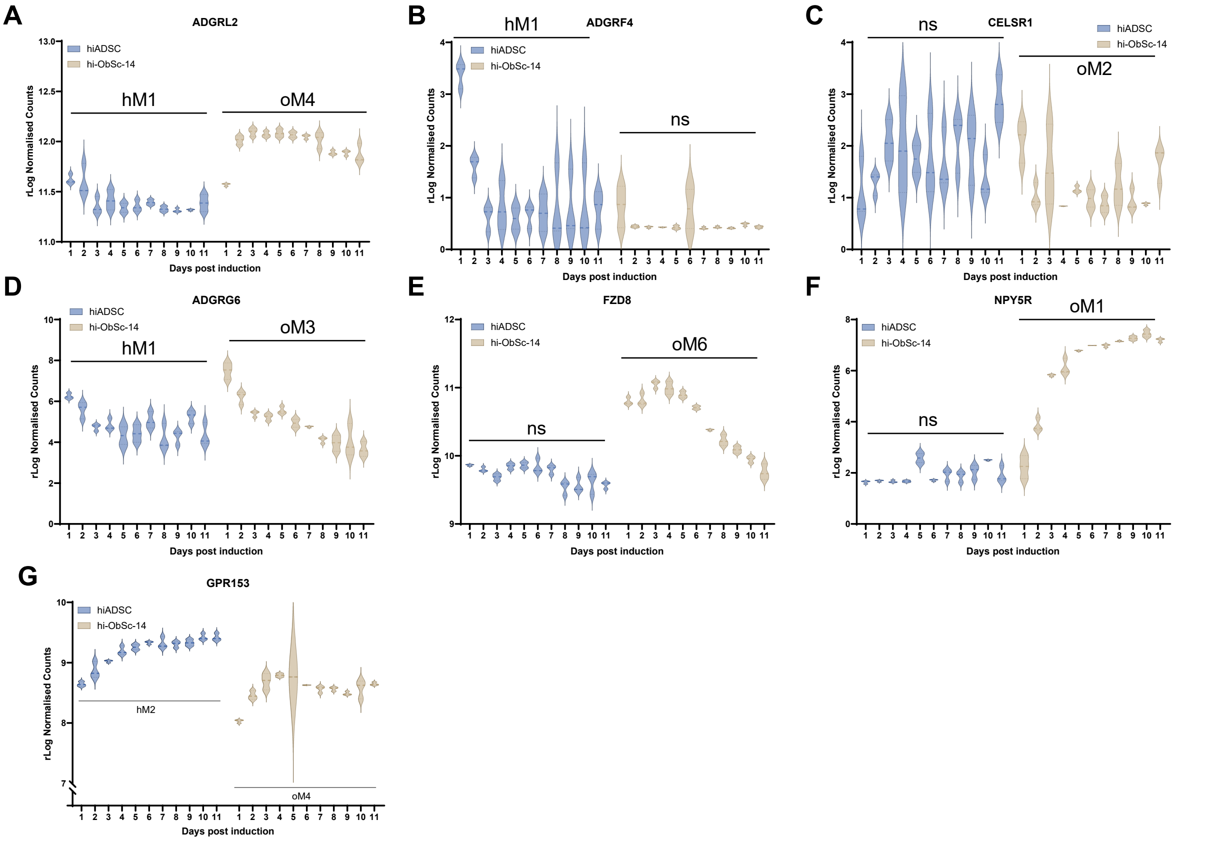


**Supplementary Figure 3: rLog Normalised counts of differentially assigned GPCRs across modules.**
**A)** Adhesion G Protein-Coupled Receptor (ADGR) L2 **B)** Adhesion G Protein-Coupled Receptor (ADGR) F4 (GPR115) **C)** Cadherin EGF LAG Seven-Pass G-Type Receptor (CELSR) 1 (ADGRC1) **D**) Adhesion G Protein-Coupled Receptor (ADGR) G6 **E)** Frizzled Class Receptor (FZD) 8 **F)** Neuropeptide Y Receptor Y5 (NPY5R). **G)** Orphan receptor GPR153. NS= not identified as significantly altered during adipogenesis defined by LRT q<0.01.

A

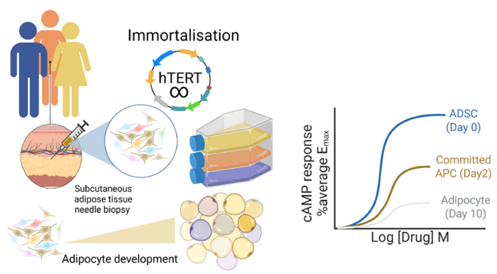


C

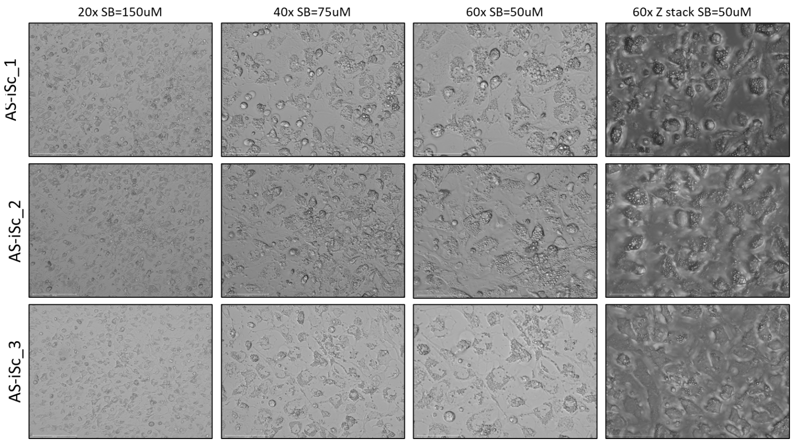


B

| Cell Line | Sex | Age |
|---|---|---|
| AS_iSc_1 | M | 35 |
| AS_iSc_2 | F | 26 |
| AS_iSc_3 | F | |
| h-iADSC (AZ) | F | 30 |
| h-iObSc-14 | F | 60 |
| h-iObSc-15A | M | 60 |
| H-iObSc-16A | F | 47 |

D

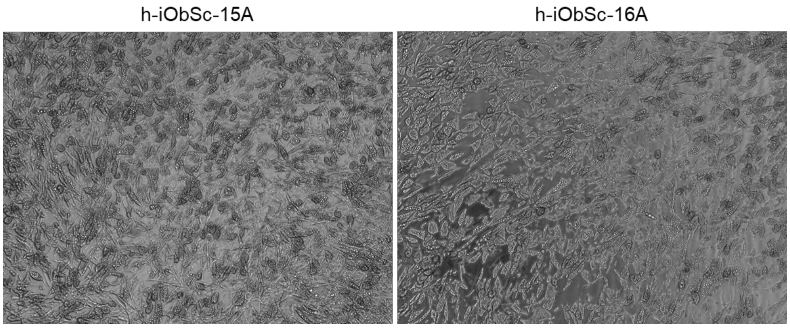


**Supplementary Figure 4: Adipocyte development in in-house derived healthy AS_iSc_1, 2 and 3 and obese h-iObSc-15 and 16. A)** Schematic overview of cell line development and experimental design. Cells isolated via needle biopsy subsequently immortalised. **B)** Donor clinical parameters for cell lines AS_iSc-1, 2 and 3. **C)** Adipogenesis in AS_iSc_1, 2 and 3 by brightfield microscopy at passage 6. Scale bar: 150, 75 and 50 μM. **D)** Adipogenesis in h-iObSc-15A and 16A (10x optical magnification) at passage 6.

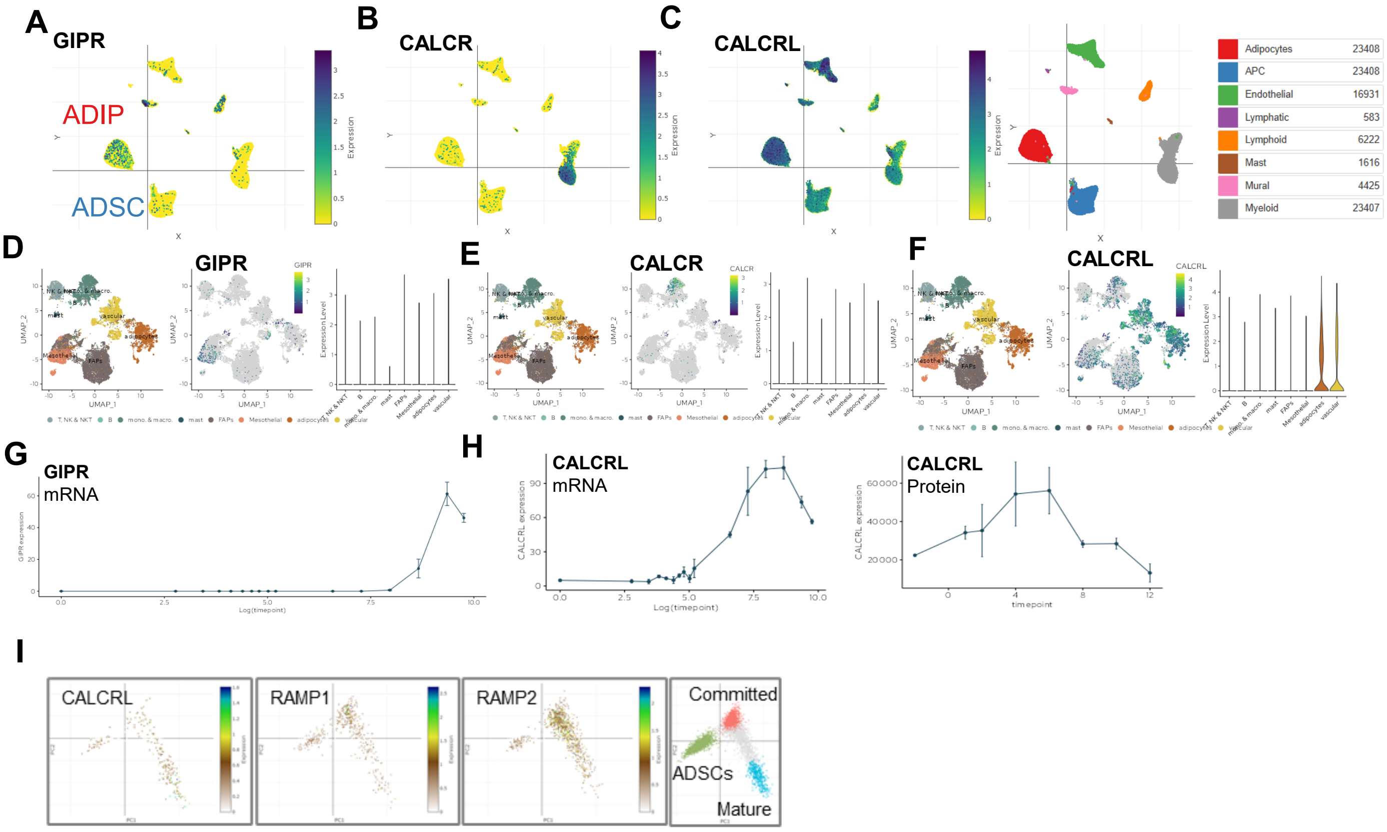


**Supplementary Figure 5: Extraction of publicly available human data sets to identify selected GPCR expression in the adipose tissue niche**

Single-nuclei RNAseq (Emont et al., 2022, Miranda et al. 2025, overlay, Single Cell Portal) **A)** GIPR **B)** CALCR and **C)** CALCRL in the adipose tissue niche. Single nuclei RNA sequencing from additional studies show **D)** GIPR **E) F) G)** Transcriptomic regulation of GIPR (source) **H)** CALCRL as induced during adipogenesis by mRNA but with alternate protein expression (intensity). GIPR and CALCR were not detected in proteomic screens. Source: Adipose Tissue Knowledge Portal (https://doi.org/10.1016/j.cmet.2025.01.012) **G) H) I)** Single-cell RNA sequencing (Loureiro *et al.,* Nat Metab 2023.) of ADSCs, committed (MGP+) adipocyte precursors and mature adipocytes identifying CALCRL, RAMP1 and RAMP2 (no detection of CALCR/GIPR or RAMP3). N= 2 independent donors. **J)** Dot plot showing the top 2 GPCRs from each 'healthy' module identified by Loureiro *et al*. Neither GIPR or CALCR were detected by this study. N=2 independent donors.
